# Phase separated ribosome nascent chain complexes paused in translation are capable to continue expression of proteins playing role in genotoxic stress response upon DNA damage

**DOI:** 10.1101/2022.03.16.484567

**Authors:** Orsolya Németh-Szatmári, Ádám Györkei, Dániel Varga, Bálint Barna H. Kovács, Nóra Igaz, Kristóf Német, Nikolett Bagi, Bence Nagy-Mikó, Dóra Balogh, Zsolt Rázga, Miklós Erdélyi, Balázs Papp, Mónika Kiricsi, András Blastyák, Martine A. Collart, Imre M. Boros, Zoltán Villányi

**Affiliations:** Department of Biochemistry and Molecular Biology, University of Szeged, Szeged, Hungary; Synthetic and Systems Biology Unit, Institute of Biochemistry, Biological Research Centre, Szeged, Hungary; Department of Optics and Quantum Electronics, University of Szeged, Szeged, Hungary; Department of Pathology, Faculty of Medicine, University of Szeged, Szeged, Hungary; Institute of Genetics, Biological Research Centre, Szeged, Hungary; Department of Microbiology and Molecular Medicine, Institute of Genetics and Genomics Geneva, Faculty of Medicine, University of Geneva, Switzerland

## Abstract

EDTA- and RNase-resistant ribonucleoprotein complexes of arrested ribosomes with protruding nascent polypeptide chains have recently been described in yeast and human cells. These complexes have been termed assemblysomes, a type of soluble condensates distinct from other known granules. Here, we use bioinformatics to identify additional proteins that likely form assemblysomes during translation. We characterize soluble condensates of the DNA helicase Sgs1, one such identified protein and a key player in the repair of DNA double-strand breaks in yeast. We show that paused ribosome-associated nascent chains of Sgs1 in condensates are able to resume translation upon UV irradiation, consistent with the return of mRNA to the ribosome pool. By extending our studies to human cell lines, we found that EDTA-resistant pellets of ribosomes from the human prostate cancer cell line DU145 are sensitive to treatment with 1,6-hexanediol, which is known to dissolve liquid-liquid phase-separated condensates. In addition, transmission electron microscopy shows that 1,6-hexanediol dissolves ring ribosomal structures from the cytoplasm of radioresistant A549 cells while making the cells more sensitive to X-rays. These results suggest that the stress response is based on a conserved mechanism involving the regulated return of phase-separated paused ribosome-nascent chain complexes to translating ribosomes.

## Introduction

Assembly of protein complexes in the dense eukaryotic cytoplasm can be challenging. Co-translational assembly of proteins can facilitate the assembly process and is essential in specific cases. Examples of this process were reported in the last few years, such as the synthesis of two adjacent proteins within the proteasome base, Rpt1 and Rpt2. While these proteins interact in their native context, they do not interact in the yeast two-hybrid assay and are not soluble when produced separately (Barrault et al., 2012; Fu et al., 2001). Instead, Rpt1 and Rpt2 are produced with ribosome pausing within EDTA and RNase resistant granules that contain the largest subunit of the Ccr4-Not complex, Not1 (Panasenko et al., 2019). These granules, referred to as Not1 containing assemblysomes (NCAs), are distinct from other known granules, such as processing bodies (P-bodies) and stress granules (SGs), as evidenced with fluorescent microscopy (Panasenko et al., 2019). P-bodies contain components of the mRNA decay machinery and are RNase- and cycloheximide-(CHX) sensitive (Youn et al., 2019; Hubstenberger et al., 2017). Stress granules contain translation initiation components and their size is between 0.1-1μm. Both P-bodies and SGs are dense entities that can be sedimented with moderate speed centrifugation (8-10000 x g) (Hubstenberger et al., 2017; Jain et al., 2016). NCAs are distinct from both of these granules as they contain ribosomes paused in translation with protruding nascent polypeptide chains, can be sedimented only by high-speed ultracentrifugation, and are resistant to EDTA and CHX treatment (Panasenko et al., 2019; Youn et al., 2019; Hubstenberger et al., 2017; Jain et al., 2016). Table 1 summarizes the basic attributes of SGs, P-bodies and NCAs.

**Table 1.** Basic attributes of stress granules, P-bodies and NCAs.

|  | Stress granules | P-bodies | NCA's |
| --- | --- | --- | --- |
| Liquid-liquid phase separation is involved in their formation | YES (Wheeler et al., 2016) | YES (Luo et al., 2018) | YES (this work) |
| Triggers in their formation | Stress induced (Wheeler et al., 2016, Kedersha and Anderson, 2002) | Constitutive in some cell lines but increase in size and number in response to stress (Wheeler et al., 2016; Kedersha et al., 2005; Ohn et al., 2008) | Constitutive in the case of DNA repair proteins (this work)<br>Induced in response to proteotoxic stress in the case of Rpt1 and Rpt2. (Panassenko et al., 2019) |
| Effect of UV | Triggers formation (Kwon et al., 2007; Moutaoufik et al., 2014 ) | No effect (Riggs et al., 2020) | Dissolves (this work) |
| Size | 100-2000 nm (Gilks et al., 2004) | 400-500 nm (Ayache et al., 2015) | 100-200 nm (this work) |
| Cycloheximide sensitivity | YES (Mollet et al., 2008) | YES (Sheth and Parker, 2003) | NO (Panassenko et al., 2019) |
| RNase sensitivity | NO (Jain et al., 2016) | YES (Sheth and Parker, 2003) | NO (Panassenko et al., 2019) |
| Markers | translation factors (such as eIF3b, eIF4A, eIF4G) and RNA-BPs such as PABP and G3BP, proteins of the small ribosome subunit (Youn et al., 2019) | components of the cytoplasmic RNA degradation machinery such as Dcp2, Dcp1 or Hedls (Youn et al., 2019) | Not1, proteins of the large ribosomal subunit (Panassenko et al., 2019, this work) |
| Velocity used for their sedimentation | 8000xg (Jain et al., 2016) | 8000xg (Hubstenberger et al., 2017) | 163 000xg (Panassenko et al., 2019, this work) |

Ribosome profiling indicated ribosome pause sites on the *RPT1* and *RPT2* mRNAs and it was determined that their translation in NCAs resumes only when the nascent chains of the partners interact (Panasenko et al., 2019). The N-terminal domains of both Rpt1 and Rpt2 are suggested to protrude out of the ribosome exit tunnel in the context of stalled translation and include the interacting helices of the two proteins. Both N-terminal protruding domains contain additionally disordered regions, a feature important for the accumulation of the paused ribosome-nascent chain complexes (RNCs) in assemblysomes (Panasenko et al., 2019). While paused ribosomes can provoke ribosome collisions and risk elimination by the ribosome quality control (RQC) mechanism, stalled Rpt1 and Rpt2 are very stable indicating that NCAs protect the paused ribosomes from RQC (Inada, 2013). This remarkable feature of NCAs most likely contributes to promoting co-translational interaction of the disordered regions of the partner proteins.

The structure of NCAs, and that how widespread they are, is not known. What we know so far about NCAs is limited to their discovery in the context of the co-translational assembly of Rpt1 and Rpt2 (Panasenko et al., 2019). A recent study using ribosome profiling revealed the pivotal role of CNOT1 in human cells in regulating ribosome pausing on numerous mRNAs, indicating that the NCA regulation might be widespread (Gillen et al., 2021). However, mRNAs translated in assemblysomes may not all show detectable ribosome pausing by ribosome profiling due to the fact that assemblysomes are RNase resistant (Panasenko et al., 2019). Given such a limitation we tried an independent approach, wherein we exploited known features of NCAs to conduct a global *in silico* analysis to identify subunits of protein complexes that may fall under NCA regulation. Using one such identified protein, the yeast Sgs1 helicase implicated in DNA damage response, we confirm that it does indeed form NCAs and that perturbation of phase separation aggravates sensitivity to DNA damage. By using a number of biochemical and microscopic approaches we demonstrate the high similarity of stalled Sgs1-RNA-ribosome complexes to Rpt1-Rpt2 containing assemblysomes and their distinctness from P-bodies and SGs. Finally, using human cell lines of tumorous origin, we present pieces of evidence to show that the role of assemblysomes in the DNA-damage response is most likely a conserved phenomenon.

## Materials and Methods

### In silico identification and analysis of protein complexes

The list of protein complexes was downloaded from the Uniprot database along with the protein and nucleotide sequences. Protein disorder data was retrieved from the MobiDB (Piovesan and Tosatto;2018) database and three different predictors were used: PONDR vsl2b (Peng et al., 2006), iUpred (Dosztányi et al., 2005a) long disorder and Espritz-Xray (Walsh et al., 2012), from which the disorder content up to the first 50 amino acids was retrieved. N-terminal lysine content and codon pairs that significantly slow down or stall translation (Ghoneim et al., 2018) were identified from the protein and the nucleotide sequences respectively, using an in-house Perl script. Amino acid changes that disrupt N-terminal protein disorder and interaction propensity were calculated using the iUpred algorithm iteratively. The algorithm was embedded in a Perl script, in each step all possible amino acid changes in the first 50 amino acids were calculated, the change was applied to the protein sequence and the process was repeated.

### Yeast strains and culture conditions

All yeast strains used for this work are listed in Supplementary Table 1. Wild type yeast cells were grown at 30°C in yeast extract peptone dextrose (YPD), while yeast cells transformed with plasmids containing genes under the regulation of copper inducible *CUP1* promoter were grown in synthetic dropout (SD) medium to exponential phase and induced for 10 min with 0.1 mM CuSO_4_ before harvesting.

### Cell culture

A549 lung adenocarcinoma and DU145 prostate cancer cell lines were purchased from ATCC and maintained in Roswell Park Memorial Institute 1640 (RPMI) medium (Biosera, Metro Manila, Philippines) complemented with 10% Fetal Bovine Serum (FBS) (EuroClone, Pero MI, Italy), 2 mM glutamine (Sigma-Aldrich, St. Louis, MO, USA), 0.01% streptomycin and 0.006% penicillin (Biowest, Nuaille, France). Cells were cultured under standard conditions in a 37 °C incubator at 5% CO_2_ and 95% humidity. When indicated, cells were treated with 1,6-hexanediol at a final concentration of 2% v/v for 30 min.

### Cloning

All plasmid and primers used for this work are listed in Table S1. NEBuilder HiFi DNA Assembly (New England Biolabs) was used for clonings. Plasmids for stalled nascent constructs were from pOP164 described previously (Panasenko et al., 2019) and obtained by PCR with oligonucleotides specific for different ORFs and cloning by NEB HiFi. The first six lysines of the Rpt1 ORF in pOP164 (pMAC1152) were generated by PCR using an oligonucleotide bearing the mutations as a forward primer. All clones were verified by sequencing and plasmids were transformed into the BY4741 yeast strain for subsequent work.

### Polysome fractionation

Cellular extracts were fractionated in sucrose gradients as described before (Panasenko and Collart, 2012). Proteins from the fractions were analysed by Western blotting.

### In vivo ubiquitination assay

The assay was performed as previously described (Panasenko et al., 2006) with cells expressing His6-ubiquitin under the control of the *CUP1* promoter and ubiquitinated proteins were purified by nickel affinity chromatography.

### Investigation of the effect of repeated UV treatment in the presence or absence of 1,6-hexanediol

Exponentially growing liquid yeast cell cultures (0.4-1 OD_600_) in 1400 ul volume were treated in a 2 cm Petri dish. Each UV irradiation lasted 2 minutes (18 mJ/cm^2^ dose). 1,6-hexanediol (SIGMA) at a final concentration of 5% v/v in 140 μl ethanol or ethanol vehicle alone was added after the first UV treatment.

An incubation time (10 min) between two UV treatments was allowed to facilitate the formation of granules. Working in the dark during sample collection for UV treated and their control samples is a necessity to avoid the effect of the efficient visible light-inducible DNA repair system available in singlecell organisms (Kelner, 1949).

### Starvation, heat and CuSO_4_ treatments

To investigate the effect of starvation on NCA formation, 50 ml yeast cells were grown ON to 6 OD_600_ and diluted with dH_2_O to 1 OD_600_ before the cells were harvested. Logarithmic growing yeast cultures in 50 ml volume were heat-stressed at 37 °C for 1 hour. 50 ml wild type yeast cells were grown to exponential phase and treated with CuSO_4_, final concentration was 1mM.

### Lysate preparation and ribosome fractionation for mRNA, rRNA, tRNA and protein examination

1 OD_600_ unit of logarithmically growing yeast cells transformed or not with plasmids containing genes under the regulation of copper inducible *CUP1* promoter were harvested and broken with 0.5 ml of glass beads in 250 μl lysis buffer. Two different lysis buffers were used for cell lysis. To pellet ribosomes and granules we used lysis buffer (20 mM HEPES (pH 7.5), 20 mM KCl, 10 mM MgCl_2_, 1% Triton X-100, 1 mM DTT, protease inhibitor cocktail), to pellet EDTA resistant granules lysis buffer with 25mM final concentration of EDTA was used. RNase inhibitor was added to the buffers in cases when mRNA was to be followed after cell lysis. Each sample was vortexed with glass beads in appropriate lysis buffer for 15 min at 4°C. Samples were centrifuged first with a short spin to pull the lysate off the beads followed by 10 min at 8000 x g at 4°C to get rid of cell debris, nuclei, aggregated proteins and SGs. Supernatants of 100 μl were either treated or not with 25mM EDTA layered on top of 500 μl 60% sucrose cushion (20 mM HEPES (pH 7.5), 20 mM KCl, 10 mM MgCl_2_ and 60% sucrose; either with 25mM EDTA or without). Samples were ultracentrifuged for 4 hours at 50.000 rpm at 4°C in a Sorvall MX 120/150 Plus MicroUltracentrifuge (Thermo Fisher Scientific) in S55A2 rotor. Pellets from ultracentrifugation were resolubilized in 2xSDS loading buffer for protein and appropriate lysis buffer for RNA examination.

The mRNAs were extracted from different fractions were analyzed by quantitative real-time PCR. For RNA extraction to examine the different rRNA and tRNA species of total extract, SG enriched, supernatant and pellet samples on non-denaturing 1% agarose gel we extracted RNA with TRI reagent^®^ (ZYMO Research) according to the manufacturer’s instructions. Proteins from different fractions were analyzed by Western blot and visualized by Coomassie Brilliant Blue staining.

In the case of human cell lines, 60% confluent cultures were harvested and treated either with 1,6-hexanediol 2% v/v, or without. Cell lysis was performed with 15 min incubation on ice in lysis buffer (100 mM KCl, 50 mM Tris-Cl (pH 7.4), 1.5 mM MgCl_2_ 1 mM DTT, 1.5% NP-40, protease inhibitor cocktail, with or without 25 mM EDTA as indicated). Ultracentrifugation and downstream analysis were similar as above using the following buffer for sucrose cushion: 100 mM KCl, 50 mM Tris-Cl (pH 7.4), 1.5 mM MgCl_2_ and 60% sucrose either with or without 25 mM EDTA. For RNA extraction to examine the different rRNA and tRNA species of total extract, SG enriched, supernatant and pellet samples on non-denaturing 1% agarose gel we extracted RNA with TRI reagent® (ZYMO Research) according to the manufacturer’s instructions. We solubilized and extracted RNA from the entire pellets (SG, pellet, EDTA-pellet), and from 500 μl of supernatants and 100 μl total extracts. We loaded half of the full volume of the RNA extracted this way on the gel from each sample.

### RNA Extraction and Quantitative Real-time PCR

After the ultracentrifugation step of yeast total protein extracts on 60% sucrose cushion with or without EDTA, total RNA was isolated from extracts obtained from different fractions using NucleoSpin TriPrep Kit (Macherey-Nagel) following the requirements of the manufacturer. RNA concentration was measured by NanoDrop spectrophotometer. Equal volumes from each fraction containing 0,1 ng - 5 μg of total RNA was reverse transcribed with oligo(dT) primers using the RevertAid First Strand cDNA Synthesis Kit (Thermo Fisher Scientific) according to the manufacturer’s instructions. Diluted cDNAs were mixed with GoTaq qPCR Master Mix (Promega) and analysed with Piko-Real 96 Real-Time PCR System (Thermo Fisher Scientific). Gene specific primers were used to detect *SGS1, RAD10, RAD14, RMI1, TOP3, RPT1, RPT2* and *ACT1*. Primer sequences are listed in Table S1.

### Western blot

Total protein samples were separated with 10% Tris-Tricine-SDS-PAGE (Schägger, 2006) then transferred to nitrocellulose membrane. After washing the nitrocellulose membranes were blocked for 30 minutes with 5% milk in TBS-T buffer (0.05% Tween 20 (pH 7.5), 150 mM NaCl and 20 mM Tris-HCl) or for 1 hour with 5% BSA (SIGMA) in TBS-T buffer. The membranes were incubated with primary antibodies overnight, at 4°C. Primary antibodies were diluted (1:1000) in milk-TBS-T or BSA-TBS-T. After several washing steps, the membranes were incubated with secondary antibodies for 2 hours at room temperature. Eventually, membranes were mixed with ECL reagent (Millipore) and signals were detected by a LI-COR C-DiGit blot scanner. The following primary antibodies were used for Western blot experiments: ANTI-FLAG M2 antibody (F1804 Sigma-Aldrich); Anti-RPS6 antibody (ab40820); Anti-RPS6 (2217 Cell Signaling Technologies); Anti-RPL11 (PA5-27468) (Thermo Fisher Scientific); Anti-Pol II (CTD4H8) (Santa Cruz Biotechnology). Goat Anti-Rabbit and Rabbit Anti-Mouse horseradish peroxidase-(HRP) conjugated secondary antibodies (Dako) were used in our experiments.

### dSTORM measurements

Super-resolution dSTORM measurements were performed on a custom-made inverted microscope based on a Nikon Eclipse Ti-E frame. EPI-fluorescence illumination was applied at an excitation wavelength of 647 nm (2RU-VFL-P-300-647-B1, P max = 300 mW, MPB Communications Ltd). The laser intensity was set to 2-4 kW/cm^2^ on the sample plane and controlled via an acousto-optic tunable filter. An additional laser (405 nm, P max = 60 mW; Nichia) was used for reactivation. A filter set from Semrock (Di03-R405/488/561/635-t1-25×36 BrightLine® quad-edge super-resolution/TIRF dichroic beamsplitter and FF01-446/523/600/677-25 BrightLine® quad-band bandpass filter and an additional AHF 690/70 H emission filter) was inserted into the microscope to spectrally separate the excitation and emission lights. The images of individual fluorescent dye molecules were captured by an Andor iXon3 897 BV EMCCD camera (512 × 512 pixels with 16 μm pixel size) with the following acquisition parameters: exposure time=25 ms; EM gain=100; temperature=-75 °C. Typically 20,000-30,000 frames were captured from a single ROI. During the measurement the Nikon Perfect Focus System kept the sample in focus. High-resolution images were reconstructed with rainstorm localization software (Rees et al., 2013). The astigmatic 3D method was applied to determine the axial position of the fluorescent dye molecules. In this arrangement, a cylindrical lens (f=4000mm) placed into the detector path introduces astigmatism, and the ellipticity value of the distorted PSFs provides information for the generation of 3D dSTORM images (Huang et al., 2008). Mechanical drift introduced by either the mechanical movement or thermal effects were analysed and reduced by employing an autocorrelation-based blind drift correction algorithm. dSTORM experiments were conducted in a GLOX switching buffer (van de Linde et al., 2011) and the sample was mounted onto a microscope slide. The imaging buffer is an aqueous solution diluted in PBS containing an enzymatic oxygen scavenging system GluOx (2000 U ml-1 glucose-oxidase (Sigma Aldrich), 40 000 U ml-1 catalase (Sigma Aldrich), 25 mM potassium chloride (Sigma Aldrich), 22 mM tris(hydroxymethyl)aminomethane (Sigma-Aldrich), 4 mM tris (2-carboxyethyl) phosphine (TCEP) (Sigma-Aldrich)) with 4% (w/v) glucose (Sigma Aldrich) and 100 mM β-mercaptoethylamine (MEA) (Sigma-Aldrich). The final pH was set to 7.4.

### Cluster analysis of super-resolved images

A density-based spatial cluster analysis (DBSCAN) was used for cluster recognition. This algorithm requires two input parameters: a minimum number of points that form a cluster (N_core_) and the maximum distance between two adjacent points (ε) (Ester et al., 1996). N_core_ and ε were set to 8 and 24 nm during the calculations which was optimal for the separation of adjacent RNCs. Quantitative characteristics of RNC clusters, such as their area or the distance of the closest neighbouring clusters (NND) were evaluated using in-built Matlab functions (convhull and pdist2). The epitope number was also estimated by a DBSCAN-based method described in a previous publication (Varga et al., 2019).

### Transmission electron microscopy (TEM)

For TEM imaging 10^5^ A549 cells were seeded onto 0.4□μm pore polyester membrane inserts (Corning) placed in a 6-well plate. Cells were left to grow until the following day when they were treated with 1,6-hexanediol in 0.5%v/v final concentration and carefully washed and fixed in 4% glutaraldehyde in PBS for 2□hours and subsequently embedded in gelatine (2% gelatine in PBS). The obtained specimen was sliced to 1-2□mm cubes, which were further embedded in epoxy (Epon 812, EMS, PA 19440) by a routine TEM sample preparation protocol. Semi-thin sections of 1□μm were prepared to identify the cell monolayer. Blocks were trimmed, thin sections of 70□nm were obtained and stained with uranyl and lead solutions. Three independent measurements where ribosomes in ring orientation were counted on at least 1500 μm^3^ cell space for each condition analyzing at least 3 cells/biological replicate were tested. Images were captured by a Jeol 1400 plus electron microscope.

### Colony forming assay

6 × 10^5^ cells/flask were seeded into T25 cell culture flasks (Biologix, Jinan, Shandong, China) and left to grow for 24 h. After 24 h of growth, the cells were either left untreated or were treated or not with 1,6-hexanediol in 0.5%v/v final concentration and were exposed to 1 Gy irradiation delivered with a Primus linear accelerator (Siemens Healthcare GmbH, Erlangen, Germany). After 30 min incubation another 1 Gy irradiation was applied on a subset of samples. On the next day cells were trypsinized, suspended in medium and counted. From each sample 700 cells/well were seeded into 6-well plates in 3 replicates and left to grow for 1 week. Then colonies were fixed in 70% methanol and 30% acetone solution and stained with 0.5% crystal violet dissolved in 25% methanol.

## Results

### Ubiquitylation is important for assemblysome formation

To understand the basic principles of assemblysome formation, we took advantage of the published construct that models the stalling of Rpt1 and the formation of assemblysomes. This construct consists of a copper-inducible *CUP1* promoter followed by sequences encoding a Flag tag, the N-terminal 135 amino acids of Rpt1, and then 12 lysine codons that lead to ribosome stalling (Figure 1A) (Matsuda et al., 2014). We found that the nascent chain of Rpt1 migrated in SDS-PAGE with a unique and discrete size but with a higher apparent molecular weight than expected. This shift is the result of ubiquitylation of nascent Rpt1, which was quantitatively altered in this context (Figure 1A). To clarify the role of this ubiquitylation, we mutated the first six lysine residues of Rpt1 and observed that nascent Rpt1 migrated with these mutations with a size that was approximately 8 kDa smaller, indicating that it was no longer ubiquitylated (Figure 1B). We tested the sedimentation profile of ubiquitylated and non-ubiquitylated Rpt1 for sucrose gradients. As previously determined (Panasenko et al., 2019), nascent Rpt1 was detected in the ribosome-containing fractions of the sucrose gradient, including the very heavy polysome-containing fractions. However, the non-ubiquitinated nascent Rpt1 was detected in the ribosome-free fractions of the sucrose gradient, suggesting that ubiquitination of the nascent chain was essential for association with ribosomes and presence in the heavy sedimenting particles (Figure 1C). The presence of the non-ubiquitinated nascent Rpt1 chain in the supernatant indicates abortive translation or translation through the stalling sequence to the stop codon. We hypothesized that ubiquitination counteracts such events to facilitate assemblysome formation. This idea served as the basis for searching for candidates of assemblysome formation by using bioinformatics (see below).

**Figure 1.**
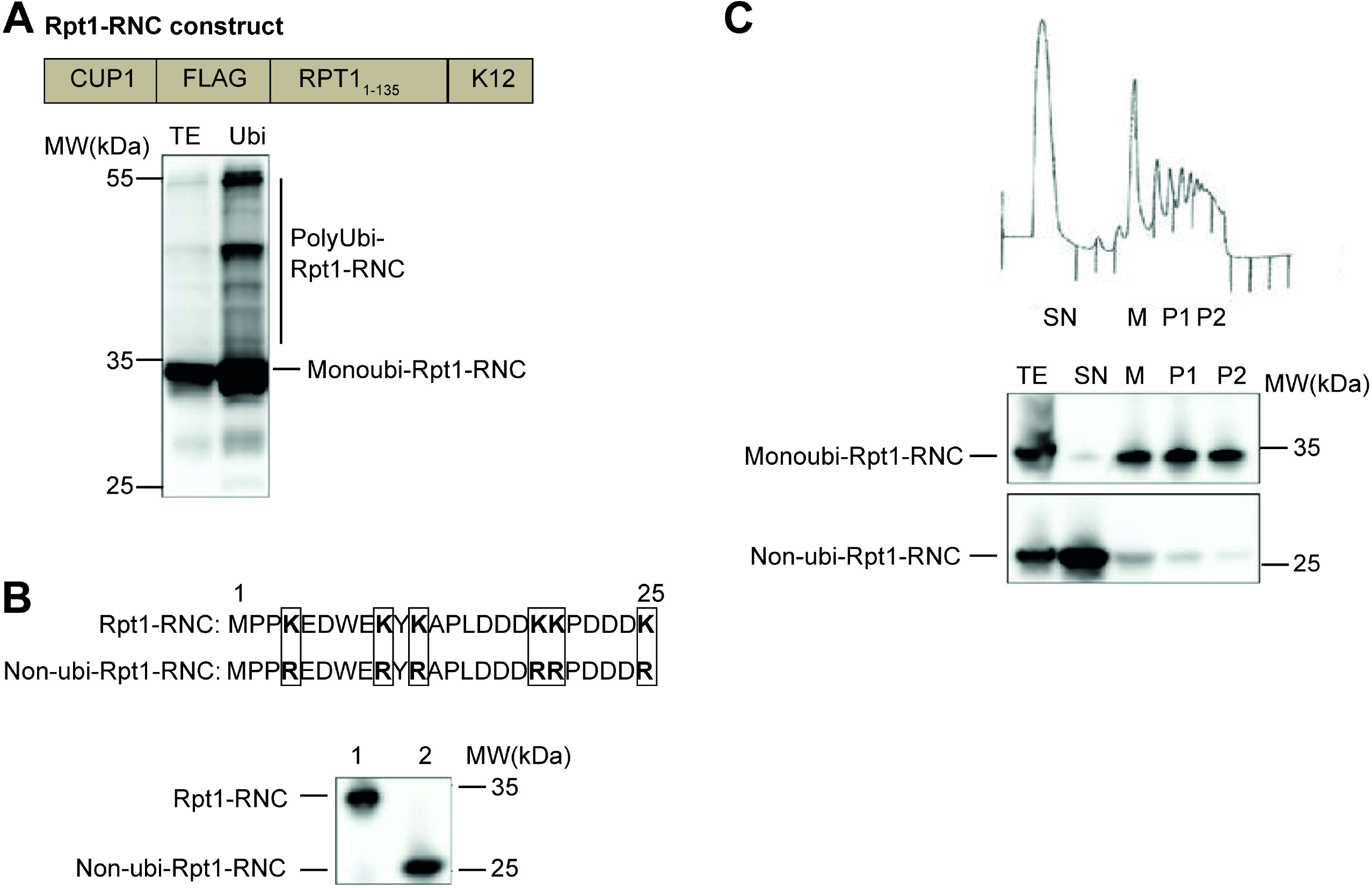
N-terminal nascent chain ubiqutination is important for NCA formation. **A:** Ubiquitinated proteins were affinity purified from cells co-expressing a stalled nascent Rpt1 chain (RNC) (RNC construct above blot) and His6-tagged ubiquitin from the *CUP1* promoter. The input extract (TE) or Ubi-affinity proteins (Ubi) were detected by Western blotting with antibodies against the N-terminal Flag tag. The discrete Rpt1-RNC is ubiquitinated (Monoubi-Rpt1-RNC). Polyubiquitinated forms of the Rpt1-RNC are additionally visible (Polyubi-Rpt1-RNC). **B:** Expression of Rpt1-RNC without (1) and with (2) the first 6 lysines mutated to arginine (mutations shown above the blot) (Non-ubi-Rpt1-RNC) monitored by Western blotting with anti-Flag antibodies. **C:** Polysome profiling of cells expressing the Monoubi-Rpt1-RNC or the Non-ubi-Rpt1-RNC (as in B). Fractions were visualized with anti-Flag antibodies. TE, total extract, SN, supernatants, M, 80S monosomes, P1 light polysomes, P2, heavy polysomes (the profile for the Ubi-Rpt1-RNC is shown above the blots with an indication of the analysed fractions respectively).

### Global scale *in silico* screen of yeast predicts DNA damage response complexes as possible assemblysome regulated candidates

Given that disordered domains within the N-terminus of Rpt1 and Rpt2 are important for NCA formation, we performed a global scale prediction of potential NCA-forming nascent proteins by ranking all known protein complex subunits of the yeast *Saccharomyces cerevisiae* according to their propensity for N-terminal disorder (Panasenko et al., 2019). We used three different assays in the *in silico* prediction of disordered N-terminal domains to generate the final ranking, but we highlight the results of a total of 6 different prediction methods (Table S2) (Peng et al., 2006; Dosztányi et al., 2005a, 2005b; Linding et al., 2003a, 2003b). The possibility of ubiquitylation of the nascent chain within the first 25 amino acids was applied as a second principle to exclude from the analysis those proteins for which the absence of lysine residues ruled them out as targets for this modification. Because stalling of the ribosome-nascent chain complexes of *RPT1* and *RPT2* mRNAs occurs at a rare codon pair, we searched for the presence of rare codon pairs among the top transcripts that came out with the previous two filters as a third filter (Ghoneim et al., 2018). Among the candidates, we identified three subunits of different complexes that play a role in the DNA damage response: Rad10, Rad14, and Sgs1 (Figure S1). Rad10 and Rad14, together with Rad1, are components of the nucleotide excision repair factor 1 (Nef1) complex. Rad14 has been shown to bind the Rad1-Rad10 nuclease to UV-induced DNA damage sites in vivo (Guzder et al., 2006). Sgs1 is part of the DNA helicase-topoisomerase complex, which together with Top3 and Rmi1 is a key player in the repair of DNA double-strand breaks (Gravel et al., 2008; Mimitou and Symington, 2008; Zhu et al., 2008).

### Certain mRNAs accumulate in assemblysomes in a manner affected by UV treatment

To support the *in silico* results, we chose to follow the mRNAs *SGS1, RAD10* and *RAD14* and tested their presence in EDTA-resistant granules. P-bodies and SGs can be sedimented by centrifugation at moderate speed (8-10000 x g), whereas ultracentrifugation through a 60% sucrose cushion of yeast extracts leads to pelleting of assemblysomes after such prior clarification of the lysate to remove P-bodies and SGs (Hubstenberger et al., 2017; Jain et al., 2016) (Figure 2A). We therefore prepared yeast extracts with or without EDTA and before and after UV treatment. EDTA dissociates the small and large subunits of the ribosome and thus destroying polysomes (Panasenko et al., 2019), whereas UV treatment induces SGs. To verify the fractionation procedure, we examined the major RNA species present in the different fractions of this separation procedure on an agarose gel. For example, the 18S rRNA, should be present in SGs containing the 40S ribosomal subunit. This was indeed confirmed in the pelleted fraction after moderate speed centrifugation (8000 x g), especially after UV treatment (Figure 2B, lane 4) (Kwon et al., 2007; Moutaoufik et al., 2014). High-speed pellets obtained in the absence of EDTA contained substantial and apparently stoichiometric amounts of 18S and 28S rRNAs along with tRNAs, as expected in polysome fractions (lanes 6 and 8). In the presence of EDTA, most of these RNAs were now detected in the supernatant (lanes 9 and 11), but detectable amounts remained in the pellet (lanes 5 and 7) suggesting the presence of ribosomes in a structure distinct from their well-known, translationally competent polysomal form, this is compatible with what we know about assembylsomes (Figure 2B, see also in Figure S2).

**Figure 2.**
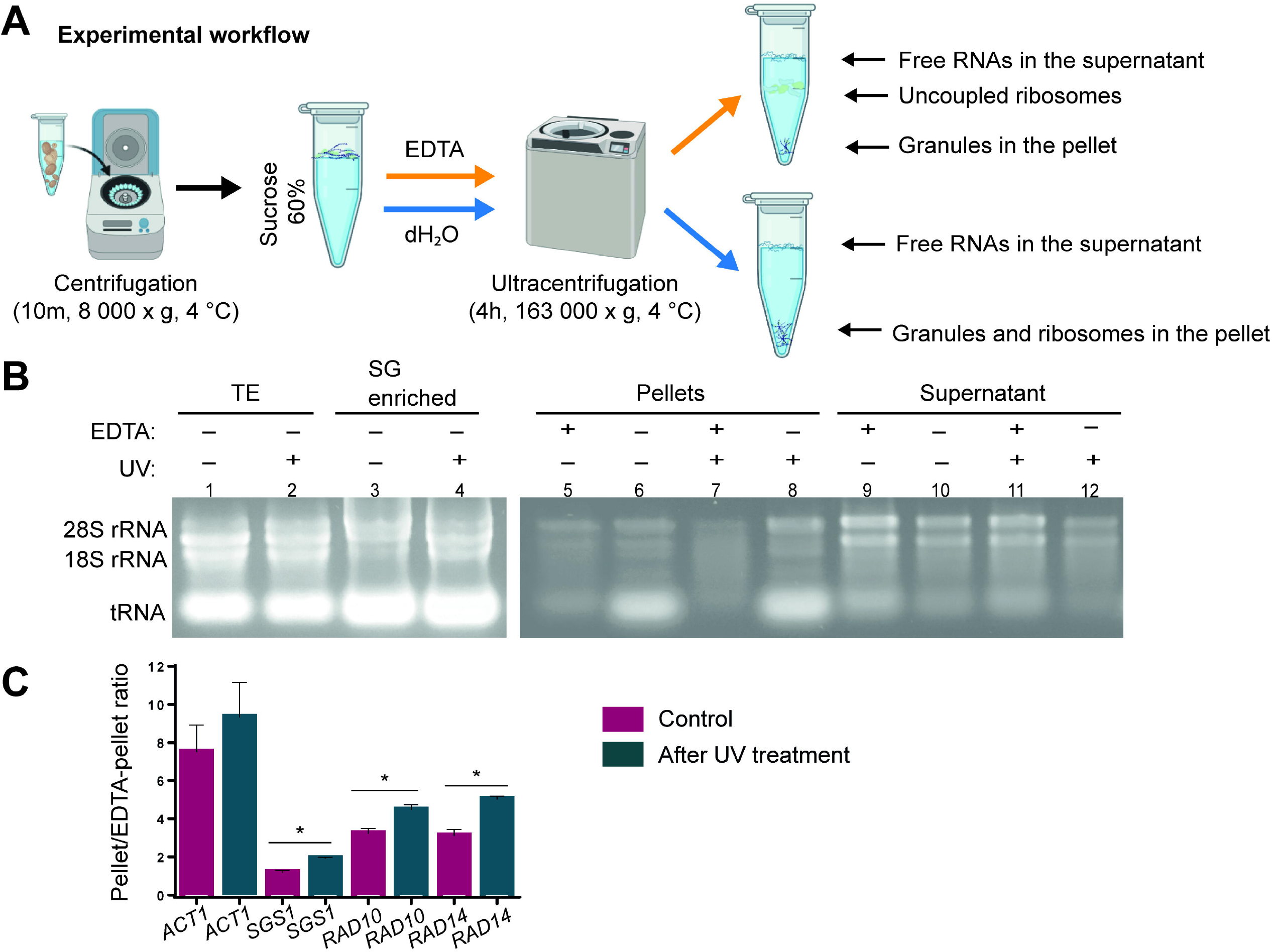
*SGS1, RAD10* and *RAD14* mRNAs are stored in EDTA-resistant, phase-separated granules. **A:** Experimental workflow for pelleting granules or granules and polysomes with ultracentrifugation of yeast total protein extracts on 60% sucrose cushions with or without EDTA respectively. Ultracentrifugation of soluble total extracts on 60% sucrose cushions leads to an enrichment of polysomes in the pellet. If the experiment is performed in the presence of EDTA the pellet contains no polysomes and is enriched in EDTA-resistant granules. SGs and P-bodies are pelleted before ultracentrifugation and EDTA treatment. **B:** Total extract of yeast cells treated with or without UV was depleted of SGs and P-bodies and separated on a 60% sucrose cushion in the presence or absence of EDTA. RNA was extracted from total extracts (TE), SG enriched, pellet, and supernatant fractions as indicated. RNA was separated on a 1% nondenaturing agarose gel. The separated RNA species are highlighted (tRNAs, 18S and 28S rRNAs). **C:** RT-qPCR analysis of mRNAs from pellets derived from identical extracts treated with or without EDTA. Results are expressed as pellet/EDTA-pellet ratio of mRNA amounts. Change in *ACT1, SGS1, RAD10* and *RAD14* pellet/EDTA-pellet ratio of mRNA quantities after the UV treatments are shown next to the same ratio measured in the untreated control. Values in the chart represent the average of two independent measurements from biological replicates, with error bars indicating standard deviation. Statistical significance was determined by a two-tailed Student’s t-test (*p < 0.05, **p <0.01).

Having established that different ribonucleoprotein entities can be separated and distinguished by our experimental procedure, we quantified specific mRNAs in high-speed pellets obtained with and without EDTA by RT-qPCR. By calculating the ratio between the mRNA content of the samples with and without EDTA, we obtain information on the amount of two different translationally engaged forms of a given mRNA, one associated with ongoing translation in “traditional” polysomes and the other in assemblysomes. mRNA species that are not translated are not considered in the subsequent calculations. According to our *in silico* prediction *ACT1* mRNA is an unlikely candidate for assemblysome formation and was selected as a control. In the absence of EDTA, approximately 8-fold more *ACT1* mRNA was detected in the high-speed pellets than in the EDTA-treated samples. This indicates that the ribosome bound *ACT1* mRNA is mostly associated with polysomes rather than with assemblysomes (Figure 2C). In contrast, the *SGS1* mRNA ratio of the -/+ EDTA samples was approximately 1, indicating a relative enrichment of *SGS1* mRNA in the EDTA fraction compared with that of the control *ACT1* mRNA. The ratio for *RAD10* and *RAD14* mRNA was approximately 3, also indicating enrichment in the EDTA fraction compared with *ACT1,* although to a lesser extent than *SGS1* mRNA (Figure 2C).

Because DNA repair proteins are required primarily in case of DNA damage, we irradiated exponentially growing wild type yeast cultures twice with UV light to mimic acute UV stress and calculated the mRNA ratios of -/+ EDTA samples as described above. Remarkably, the ratios for mRNAs *SGS1, RAD10* and *RAD14* increased after UV treatment in all cases, indicating a shift of mRNAs from EDTA-resistant to EDTA-sensitive fractions. However, there were no significant elevations in total mRNA levels (see further in Figure 3D). The most likely explanation for this observation is that translation of mRNAs in the EDTA-resistant pellets resumed, which resulted in a loss of condensation of these mRNAs in assemblysomes, to meet the requirements of the DNA damage response processes (Figure 2C).

**Figure 3.**
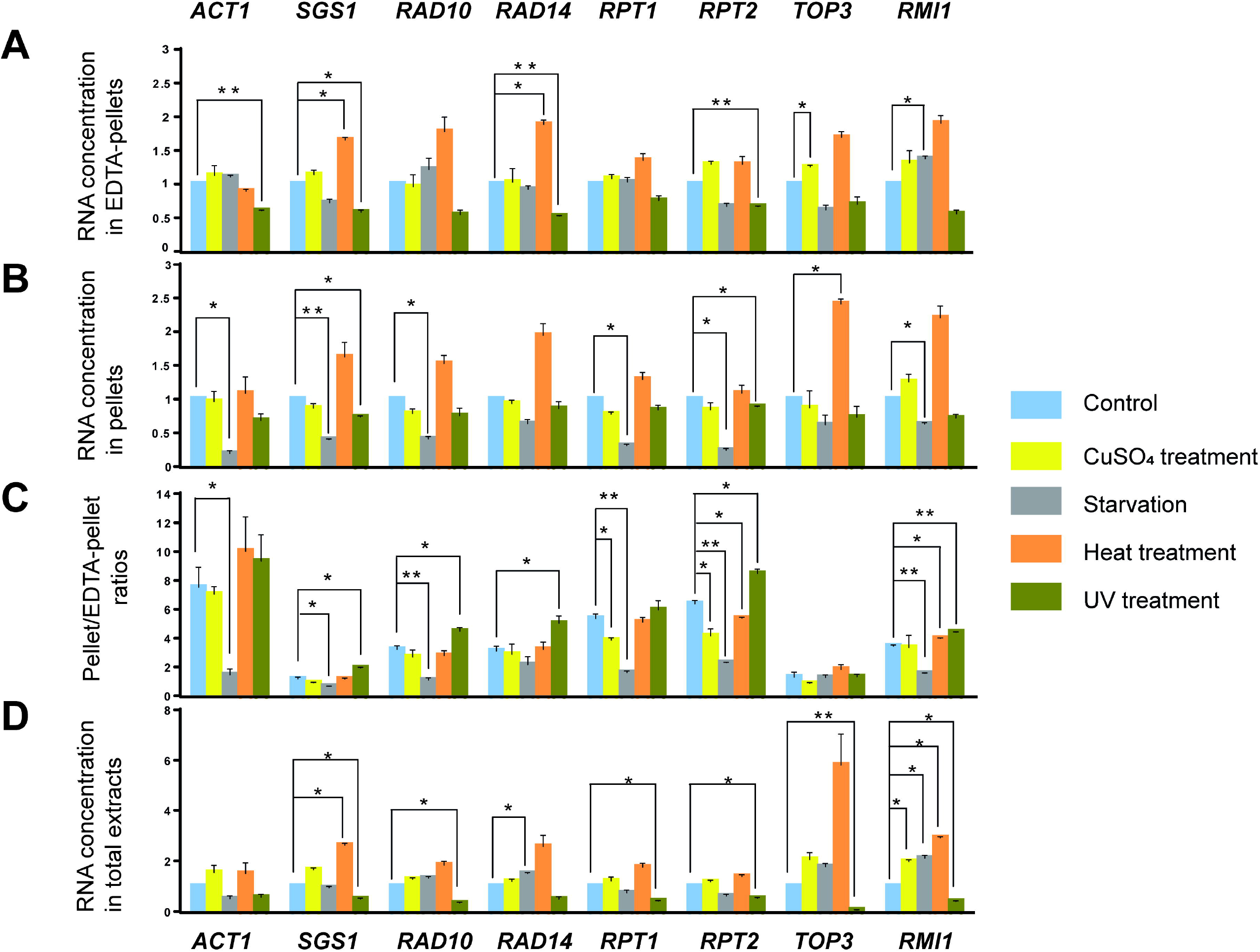
mRNA content of EDTA-pellets is stress dependent. RT-qPCR analysis of mRNAs from total extracts and pellets originating from identical extracts treated with or without EDTA. Fold change in different amounts of mRNA after the indicated treatments compared with the untreated control (blue bars set to 1) in EDTA-pellets (**A**) and in pellets (**B**). The pellet/EDTA-pellet ratio (**C**), and the fold change in total extracts (**D**) of the indicated mRNAs are also shown. The yellow, grey, orange, and green bars represent the changes measured after CuSO_4_, starvation, heat, and UV treatments, respectively. Values in the chart represent the average of two independent measurements from biological replicates, with error bars indicating standard deviation. Statistical significance was determined by a two-tailed Student’s t-test (*p < 0.05, **p <0.01).

### Genotoxic and proteotoxic stresses have opposing effects on the mRNA content of ribosomes and EDTA-resistant ribosome pellets

The above results prompted us to investigate a range of stress conditions including heat, CuSO_4_ treatment, and starvation in addition to UV irradiation (Figure 3). The mRNAs *RPT1* and *RPT2* were included in the analysis because they are known to accumulate in NCAs under proteotoxic stress such as heavy metal or heat treatment. We also included mRNAs encoding two other subunits of the Sgs1-Top3-Rmi1 complex to see if the interaction partners of Sgs1 are regulated by assemblysome formation.

Treatment with CuSO_4_ had little effect on the presence of most of the mRNAs examined in highspeed pellets (Figure 3A and B). However, there was a markedly reduced -/+ EDTA pellet ratio of *RPT1* and *RPT2* (Figure 3C), consistent with the published behavior of *RPT1* and *RPT2:* accumulation in NCAs during proteotoxic stress.

Starvation resulted in a decrease in all mRNAs in high-speed pellets in the absence of EDTA (Figure 3B), most likely due to the formation of SGs (Reineke et al., 2018) and in -/+ EDTA pellet ratios (Figure 3C), because there was no significant change in mRNAs in high-speed pellets in the presence of EDTA (Figure 3A), i.e. in assemblysomes.

In response to heat, the only apparent change was an increase in mRNA levels for some DNA repair protein-encoding mRNAs, overall (Figure 3D) and in high-speed pellets in the presence (Figure 3A) and absence (Figure 3B) of EDTA, but not significantly in all cases and with no effect on -/+ EDTA ratios (Figure 3C). Thus, heat shock increases the relative presence of DNA repair protein-encoding mRNAs in assemblysomes but has no effect on their association with EDTA-sensitive ribosomes.

The UV-induced genotoxic stress response to high-speed pellets in the presence and absence of EDTA was very different from the response to these other stresses described above. We observed a marked decrease in mRNA levels in high-speed EDTA-pellets (Figure 3A) and to a lesser extent in pellets without EDTA (Figure 3B), such that the pellet/EDTA-pellet ratio was elevated in the case of all studied mRNAs (Figure 3C). The change in the pellet/EDTA-pellet ratio compared with the untreated control was more pronounced for DNA repair protein-encoding mRNAs than for other mRNAs, with the exception of *TOP3*. For example, the pellet/EDTA-pellet ratio almost doubled in the case of *SGS1* mRNA after UV treatment compared with the control (Figure 3C).

Taken together, the above results suggest that the mRNAs in the condensates are regulated in a stress-dependent manner in unison. In response to proteotoxic stress, the distribution of mRNAs partitioning to assemblysomes increases, whereas in response to genotoxic stress, mRNA condensates released whether or not they are useful in counteracting the consequences of the stress.

### Granules contain multiple identical RNCs and their density is ORF dependent

To provide visual evidence that the nascent chain of Sgs1 is involved in the formation of assemblysomes as was described for Rpt1, we created a construct for Sgs1 similar to the one originally used to characterize the Rpt1-RNC and to demonstrate the existence of EDTA-resistant NCAs (Panasenko et al., 2019) (Figure 4A). We cloned the first 135 amino acids of the coding sequence (CDS) of *SGS1* in frame with a stalling sequence instead of the rare codon pair in the CDS of *SGS1* (Figure 4A). This happens to be identical to the length of the originally studied Rpt1-RNC (Figure 1A). To compare the granule-forming ability of these similar inducible constructs and to obtain information about the physical properties of the RNC granules in the cytoplasm, we performed dSTORM imaging on yeast cells transformed with the Rpt1-and Sgs1-RNC-expressing plasmids. The approximate size of the ribosome is 30 nm in diameter, and the resolution of dSTORM microscopy allows us to estimate (van de Linde et al., 2011; Nieuwenhuizen et al., 2015) the number of RNCs even when the ribosomes are densely packed in the granules. The dSTORM microscopy showed that expression of both constructs results in clustered staining in the cytoplasm. This indicates that the nascent chains are not present as single soluble RNCs that would stain the entire cytoplasm dispersedly but are rather present in granules with multiple RNCs inside (Figure 4B). The clustered staining of Sgs1-RNC and Rpt1-RNC was very similar and confirmed that the N-term of Sgs1 is also capable of forming condensates in the cytoplasm, similar to what has been reported for Rpt1-RNC (Panasenko et al., 2019). We noted minor differences in the number and density of granules between the two constructs (Figure 4A). The lateral size (area) of granules was larger in Sgs1-RNC expressing cells with similar median than in Rpt1-RNC (Figure 4C). The medians are identical because of the high number of individual RNCs that are not in granules (Figure 4C-4E). We did not find a typical geometric shape for a granule, but their density in the cytoplasm was different between the two construct-expressing cells, according to the average distance of the nearest granule (Figure 4E). The average epitope number, indicating the density of epitopes within granules, was also slightly different between the two constructs studied. This value ranged from an average of 5 in Rpt1-RNC granules to an average of 8 found in the Sgs1-RNC granules (Figure 4F). In parallel, these values correlated with the slight differences in lateral size of the granules in the range of several thousand square nanometers (compare Figure 4D and 4F for both constructs). The latter comparison suggests that for each epitope representing a nascent chain, the granules contain something large that is compatible with the size of the ribosomes.

**Figure 4.**
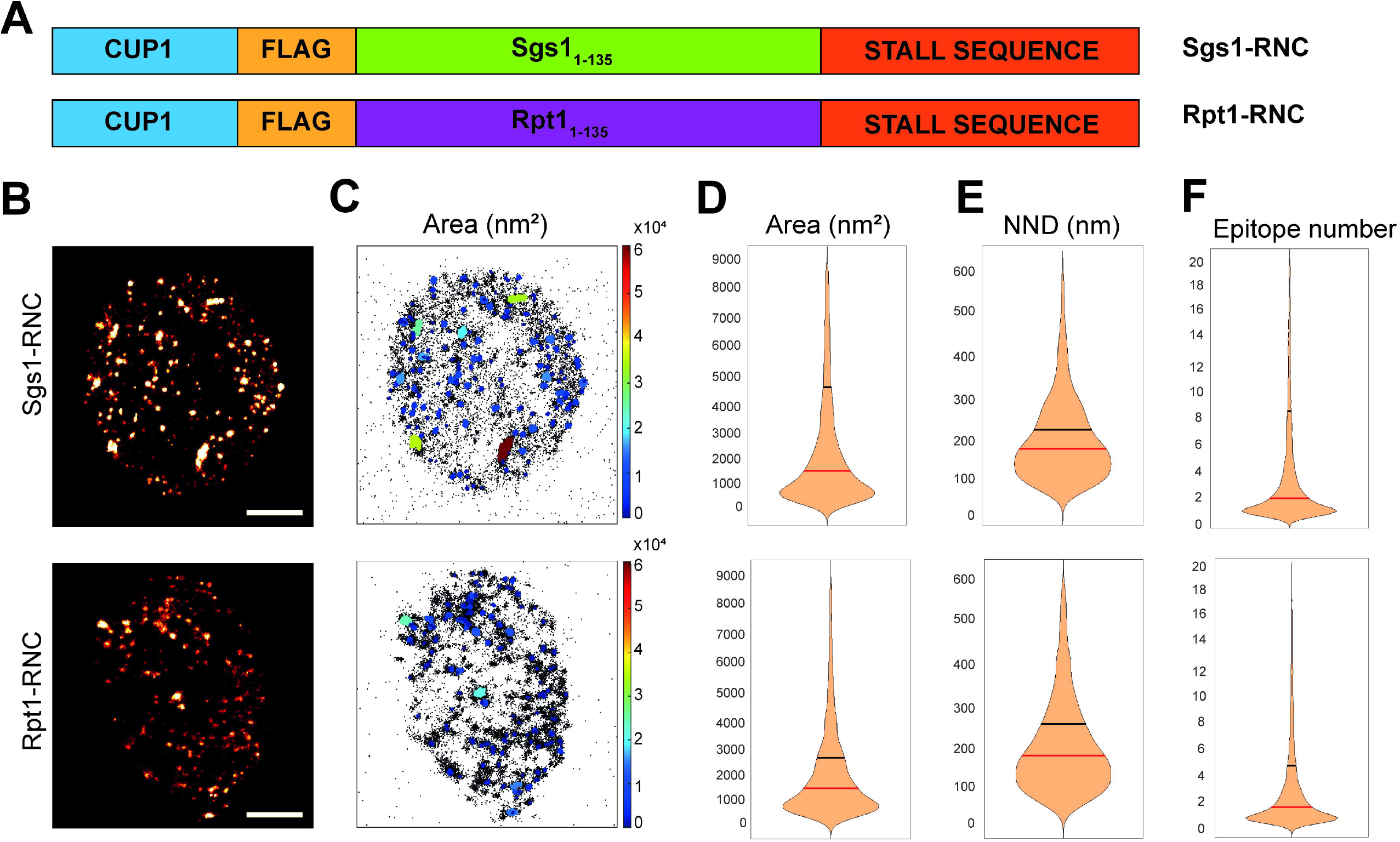
Rpt1- and Sgs1-RNCs form granules in the cytoplasm. **A:** Schematic map of the two constructs expressing either Rpt1- or Sgs1-RNC that are identical apart from their first 135 amino acid encoding sequences. **B:** Two representative reconstructed dSTORM images reveal the granular staining of N-terminally Flag tagged Rpt1-RNCs and Sgs1-RNCs. Staining was performed using M2 Flag antibody followed by secondary Alexa647 antibody. Scalebar: 1 μm. **C**: Map of the cluster analysed localizations via DBSCAN algorithm. The clusters are color-coded based on the size of their area. Increasing color temperature represent larger size granules as represented on the right. Violin plots represent the different distribution of clustered Flag signal area in nm^2^ (**D**), of nearest neighbor distance (NND) in nm (**E**), and the distribution of the epitope number/cluster (**F**), comparing 36 different Sgs1-RNC expressing cells and 68 different Rpt1-RNC expressing cells, from 2 (Rpt1-RNC) and 3 (Sgs1-RNC) independent STORM measurements done on both constructs. Vertical lines represent the median (red) and the mean (black) values.

### Nascent Sgs1 in granules are ribosome-associated

The density of Rpt1- and Sgs1-RNC granules detected by dSTORM shown on Figure 4 strongly suggests that the identified granules contain ribosomes and are therefore neither composed of released nascent chains nor irreversibly aggregated RNCs.

To gain further evidence that the Sgs1 nascent chains obtained by high-speed centrifugation in the presence of EDTA are indeed ribosome-associated, we compared the levels of the ribosomal protein Rps6 in EDTA-pellets with or without inducing expression from the Sgs1-RNC expressing plasmids. The rational of this experiment is that increasing the amount of Sgs1-RNC should enrich ribosomes proportionally. Indeed, such enrichment of ribosomal Rps6 protein was evident in both -/+ EDTA pellets when copper-induced and non-induced samples were compared (Figure S2). Remarkably, Coomassie staining also confirmed the enrichment of proteins in pellets and EDTA pellets in the copper-induced sample in the range of ribosomal proteins in size. This result is consistent with our other observations presented above and independently demonstrates that ribosomes are part of EDTA-resistant granules.

### Sgs1-RNC in pellets consists of multiple different size bands compatible with mono- and polyubiquitinated forms similar to Rpt1-RNC

Previously, it was shown that the disordered regions of the N-terminal domains of Rpt1 and Rpt2 are key to their assemblysome association (Panasenko et al., 2019). To determine whether the disordered N-terminal region of Sgs1 has a similar function in granule association, we used an algorithm and the iUpred disorder prediction method (Dosztányi et al., 2005a) to determine the minimal amino acid change that would disrupt the disordered nature of the N-terminal domain of Sgs1 (Figure 5A). We generated these mutations in the Sgs1-RNC expressing plasmid, transformed and induced the mutant and non-mutant constructs in yeast. To follow the expression of soluble and aggregated Sgs1-RNCs to be compared in the two constructs, we prepared extracts by postalkaline lysis (PAL), which is known to extract both aggregated and soluble proteins. We analysed identical OD_600_ units of cells after identical copper induction and compared the levels of nascent Sgs1 chains and Rpl11 as a loading control (Figure 5B). The mutant nascent Sgs1 was detected not only as a major form, but also in an 8 kDa smaller form, and as a smear above the main band, which is most likely due to increased polyubiquitination as was observed with Rpt1-RNC (Figure 1A, Figure 5B, C). Due to these similarities to the ubiquitylation states of Rpt1-RNC, we refer to the different forms of Sgs1-RNC with the same nomenclature (as non-, mono- or polyubiquitylated). The increased presence of polyubiquitylated Sgs1-RNC in case of the mutant suggests that the disordered structure protects Sgs1 nascent chains from RQC upon ribosome stalling (see also Figure 5D).

**Figure 5.**
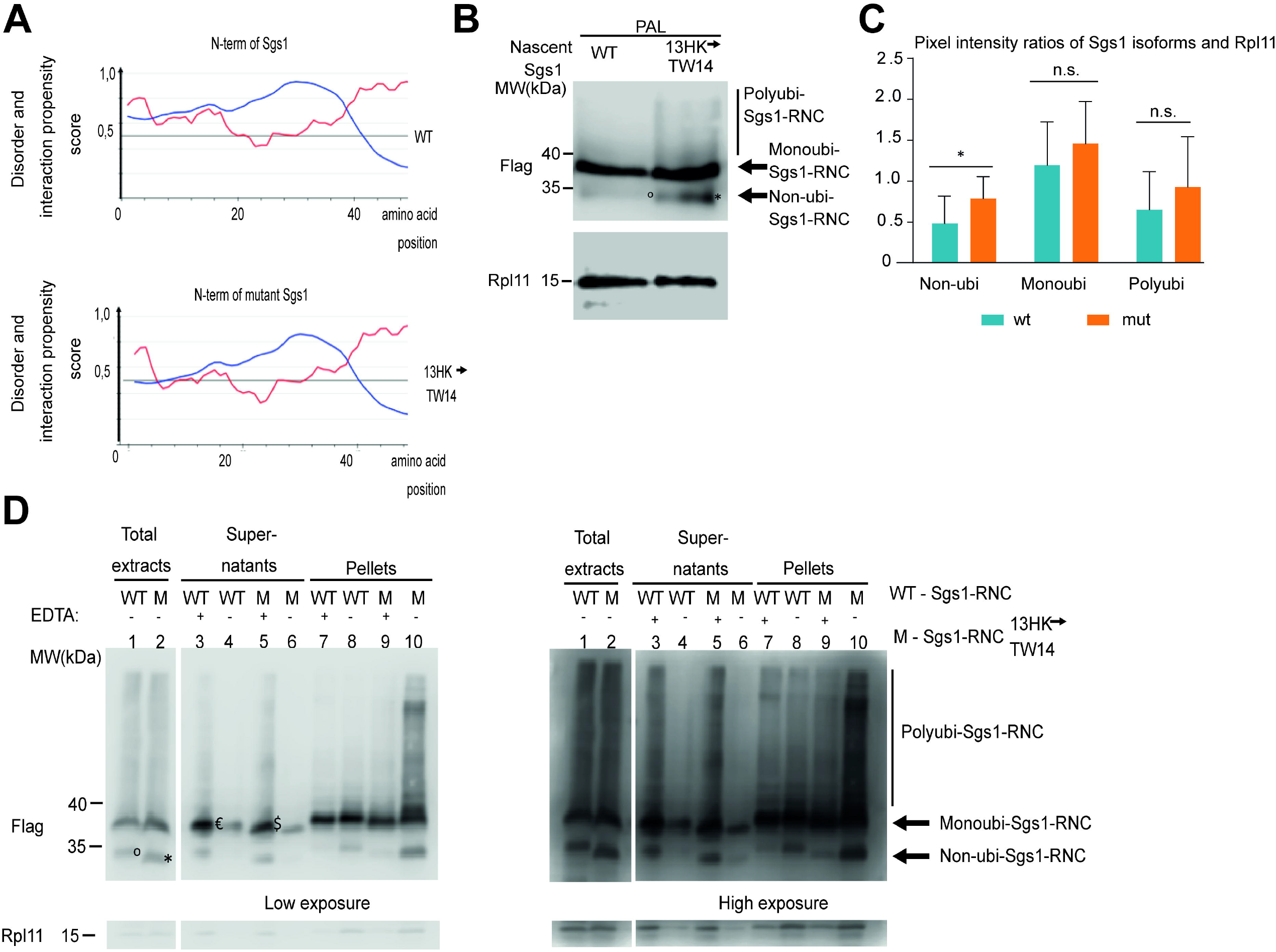
Sgs1-RNC sediments in pellets after ultracentrifugation through a 60% sucrose cushion even in the presence of EDTA. **A:** Diagrams show protein binding region and disorder propensity of the first 45 amino acids of wild type and mutant (13HK-TW14) Sgs1 as revealed by ANCHOR2 (Dosztányi et al., 2009) (blue line) and by IUPRED2 (Dosztányi et al., 2005a) (red line) algorithms respectively. **B:** Western blots show Flag-tagged wild type (WT) and mutant (13HK-TW14) Sgs1-RNCs extracted with post alkaline lysis (PAL). Rpl11 is revealed for loading control. The main band is marked with an arrow (Mono-ubi-Sgs1-RNC), as well as the 8 kDa smaller than main band (Non-ubi-Sgs1-RNC). Size difference is apparent for the Non-ubi-Sgs1-RNC between wt and mutant and marked with ° and * respectively. Polyubiquitylated Sgs1 forms are indicated (Polyubi-Sgs1-RNC). Molecular weight markers are highlighted in kDa (MW). **C:** Quantification of the specific bands (Non-ubi-Sgs1, Monoubi-Sgs1, Polyubi-Sgs1) as shown in B using three biologically independent Western blots performed with Flag tagged mutant (13HK-TW14) and wild type (WT) Sgs1-RNC’s extracted with post alkaline lysis (PAL). Pixel intensities were determined by the ImageJ software for these bands and also for Rpl11 and the ratio of these measurements are shown to correct for loading. Standard deviation and statistical significance of differences between the expression of specific bands between wt and mutant are shown. Statistical significance is determined by paired Student’s t-test (*p < 0.024) **D:** Copper-induced Flag-Sgs1-RNC and Flag-Sgs1 (13HK-TW14) derivative expressing yeasts were extracted: Total extracts, pellets and supernatant fractions were analyzed - after ultracentrifugation through a 60% sucrose cushion - by 10% SDS PAGE separation followed by Western blotting with Flag antibodies. Extracts were either untreated or treated with 25 mM EDTA before loading onto the sucrose cushion (EDTA). The arrows indicate an 8 kDa faster migrating non-ubiquitinated protein, which is also marked with ° in the case of wt and * in the case of the mutant. The mono-ubiquitinated form of Flag-Sgs1 is highlighted with € and $ in wt and the mutant, respectively. The polyubiquitinated Flag-Sgs1 ladder is also highlighted. These bands are reminiscent of similar results obtained with Rpt1-RNC in Fig. 1. As a loading control, we show the Rpl11 revelation. For clarity both low and high exposure revelations of the Western blots are shown.

We examined the behavior of the two Sgs1-RNC expressing constructs in ultracentrifugation experiments performed through 60% sucrose cushions (Figure 5D). In these experiments, we collected and analyzed two different fractions, the high-speed pellet and supernatant samples (Figure 5D). The different expression patterns of Sgs1 with or without the mutation were less pronounced in the soluble extracts (compare the lanes in Figure 5B with the first two lanes in Figure 5D). The Sgs1-RNC behaved like the Rpt1-RNC and was present in the pellet after ultracentrifugation through a 60% sucrose cushion even in the presence of EDTA (Figure 5D, lane 7). To show that this was not simply due to overexpression of a protein by the *CUP1* promoter, we cloned Rpb9 into the same plasmid with additional stop codon upstream of the truncation sequence as a control (Figure S3A). Rpb9 is a part of RNA polymerase II (RNAPII), which has 12 subunits in yeast. It has been reported that Rpb9 is the last subunit to assemble into the RNAPII complex, and importantly, that this association is post-translational (Acker et al., 1997). Rpb9 did not accumulate in polysome fractions or in EDTA-pellets. It was released from ribosomes and was detectable in the supernatant as expected (Figure S3B).

In addition to the main form of Flag-Sgs1-RNC, which migrates around 38 kDa (Fig. 5D, shown with the arrow “Monoubi-Sgs1-RNC”), we detected a faster migrating form, about 8 kDa smaller, in the total extracts (Figure 5D, lane 1 marked with an °) also mainly in the pellets obtained without EDTA (shown with the arrow “Non-ubi-Sgs1-RNC” on Figure 5D, lane 8). This ~30 kDa isoform was detectable in supernatants treated with EDTA (Figure 5D, lane 3). We also detected a smear of slower migrating forms that migrated above the main 38 kDa band to the top of the gel and were more evident in the EDTA-treated supernatant than in the supernatant without EDTA (shown with a bracket “Polyubi-Sgs1-RNC” on Figure 5D, lane 3, 4). In the experiments performed with the Rpb9 expressing construct, where the protein is released from ribosomes and found only in the supernatant, this smear is absent (Figure S3). These results are consistent with polyubiquitination of Sgs1-RNC upon ribosome stalling. The 8 kDa faster migrating band appears to be ribosome bound because it is absent in the supernatant but present in the pellet without EDTA. Conversely, it is present in the supernatant but reduced in the pellet in the presence of EDTA (Figure 5D). This is reminiscent of the results obtained with Rpt1-RNC showing that the non-ubiquitinated form cannot form EDTA-resistant condensates that sediment in heavy sucrose fractions (Figure 1C).

Experiments performed with the Sgs1-RNC non-disordered mutant yielded very similar results to those described above for the wild type Sgs1-RNC. However, the protein expressed from the same construct, differing only by two amino acid substitutions, appeared slightly smaller than the 38 kDa major form of wild type Sgs1-RNC (Figure 5D, compare the major bands in lanes 3 and 5, marked with € and $, respectively). Interestingly, the polyubiquitinated smear in the pellets of the mutant is much stronger as well as the 30 kDa band (Figure 5D, compare lanes 9, 10 to 7, 8). The amount of the main 38 kDa band in EDTA-pellets appears to be similar in both constructs.

After addition of EDTA, polyubiquitinated forms specific for mutant Sgs1 were reduced in the pellet (Figure 5D, compare smear above 38 kDa in lanes 9 and 10) and increased in the supernatant (compare lanes 5 and 6). A possible reason is that EDTA can dissolve aggregates compatible with the density of polysomes that were originally in the pellet to the supernatant. Increased aggregation of mutant Sgs1 explains the discrepancy between the comparison of samples prepared after alkaline lysis (Figure 5B) and soluble extracts (lanes 1, 2 in Figure 5D). The nearly identical TE samples that served as input for the ultracentrifugation experiments do not explain the difference visible in the pellets extracted from the two constructs. The differences in non-ubi-Sgs1-RNC and polyubi-Sgs1-RNC are negligible between the wild type and mutant Sgs1 forms in the case of the soluble extracts, but the difference is significant at least for the non-ubi-Sgs1-RNC bands from the samples prepared with postalkaline lysis (Figure 5C). The difference in polyubiquitinated nascent Sgs1 - despite the small difference in inputs - may be due to a difference in enrichment of the pellet without EDTA between the experiments with the two constructs (Figure 5D, compare lanes 1,2 with 8,10). Different enrichment might occur due to pelleting of aggregated polyubiquitinated proteins only in the mutant.

### 1,6-hexanediol dissolves EDTA resistant ribosome assemblies

The fact that the N-terminal disordered protein structure appears to be important for the formation of condensates that protect both the protein and mRNA from degradation suggests that ribosomes may phase separate when a disordered protein protrudes from them. It has been previously reported that disordered protein structures are important for phase separation (Alberti, 2017). To address this hypothesis, we tested whether 1,6-hexanediol (HEX) resolves condensates containing *ACT1, SGS1*, and *RAD10* mRNAs. Although HEX may have a pleiotropic effect on cell physiology, it is commonly used as a compound that efficiently inhibits protein phase separation by disrupting weak hydrophobic protein-protein or protein-RNA interactions required for the formation of dynamic liquid-like assemblies (Kroschwald et al., 2017). Ultracentrifugation experiments were then performed in the presence of HEX to determine whether the mRNAs in the EDTA-pellets were of phase-separated origin in granules. After treatment with HEX, the levels of the mRNAs *SGS1* and *RAD10* in the EDTA-pellets were greatly reduced in UV-treated yeast cells, indicating that liquid-liquid phase separation is indeed involved in the formation of the granules (Figure S4A). The HEX-sensitive sedimentation of *ACT1* mRNA in the EDTA-pellets was much less obvious (Figure S4A).

To further elaborate on the phase-separated nature of EDTA-resistant condensates, we decided to switch model species and test whether HEX sensitive EDTA-resistant condensates also exists in higher eukaryotes. We prepared total extracts from DU145 prostate cancer cells and pelleted them by ultracentrifugation through a 60% sucrose cushion in the presence of EDTA. The total extracts, supernatant, TCA-precipitated sucrose cushion, and EDTA-pellets were analyzed from mock and HEX treated DU145 prostate cancer cells by tracking the ribosomal protein Rps6. The Rpb1 subunit of RNAPII was used as a loading control. Ribosome content was indeed reduced in the HEX-treated EDTA pellet (Figure S4B), suggesting that phase separation of translating ribosomes is an evolutionarily conserved phenomenon.

### Phase-separated granules are present in the cytoplasm of A549 human alveolar adenocarcinoma cells

Our results with yeast cells may represent a general phenomenon, and the formation of NCA-like phase-separated granules may be important for radiation-resistance in higher-order eukaryotes. To support this idea, we searched for a radiation-resistant cell type. The A549 lung adenocarcinoma line was selected to study densely packed ribosomal assemblies similar to those of phase-separated Rpt1- and Sgs1-RNC-containing granules of yeast cells. Ribosomes have a distinct morphology in electron micrographs (Igaz et al., 2020), so we decided to follow their higher-order arrangement with transmission electron microscopy (TEM). Electron micrographs of fixed A549 cells revealed densely packed ribosomes that were mostly arranged in a ring-like orientation (Figure 6A). These structures were approximately 100 nm in diameter, consistent with the size of 8-9 tightly packed ribosomes that could be detected with STORM by following the Flag-tagged nascent Sgs1-RNC protruding from the ribosome (Figure 4F, upper panel). To confirm that these ribosomes were components of phase-separated granules, we performed staining with A549 cells treated with 2% v/v HEX before fixation. Indeed, the presence of ring-oriented ribosomes was markedly reduced in the cytoplasm of A549 cells treated with HEX compared with untreated cells (Figure 6B and C).

**Figure 6.**
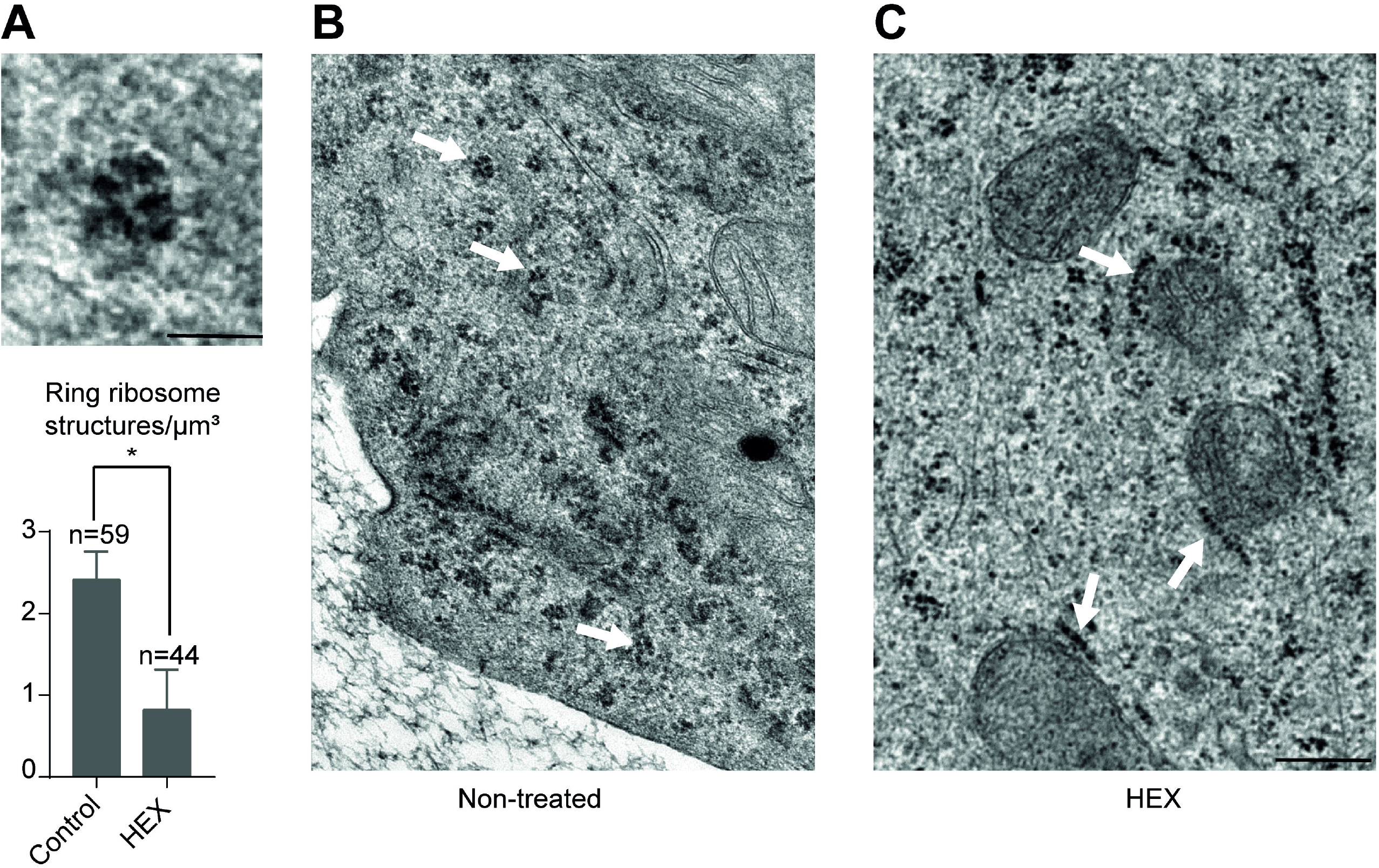
Closely packed ribosomes sensitive to 1,6-hexanediol treatment are detectable in the cytoplasm of A549 cells. **A:** Typical example of ring-like ribosome structure as revealed by TEM after fixing with glutaraldehyde (top) and diagram of quantification of combined results from three independent experiments in which ring ribosomes were counted on at least 1500 μm^3^ cell space for each condition analyzing at least 3 cells/biological replicate (with/without HEX) (bottom). Statistical significance was determined by a two tailed Student’s t-test (*p < 0.05). Scalebar: 50 nm. **B:** Cells were fixed with glutaraldehyde and examined with TEM. White arrows are indicating typical ring-like ribosome structures, the most common appearance of adjacent ribosomes in non-treated cells. **C:** Cells were treated with HEX for 1 hour and fixed with glutaraldehyde for TEM as in panel B. Same magnification as in panel B. White arrows are indicating linearly arranged ribosomes, the most common appearance of adjacent ribosomes in HEX treated cells. Scale bar: 500 nm

### 1,6-hexanediol sensitizes A549 cells to ionizing radiation

We conducted viability assays to look for the role of phase separated ring-oriented ribosome containing granules in DNA damage response. An equal number of A549 cells were treated or not with HEX, and we compared the number of surviving colonies after a single irradiation of 1 Gray dosage or after successive treatments of 1 Gray with a 30-minute incubation period between irradiations (Figure 7). While a single irradiation significantly decreased cell viability, cells treated with HEX were no more sensitive to ionizing radiation than the untreated control. The viability of consecutively irradiated cells without HEX treatment, remained comparable to that of cells after single irradiation, despite twice the radiation dose. However, the viability of cells treated with HEX after double irradiation was significantly lower than that of cells treated with HEX and irradiated only once, and also than that of consecutively irradiated cells not treated with HEX (Figure 7). These results point toward a mechanism regulated by phase separation, which is required for DNA damage response in these cells.

**Figure 7.**
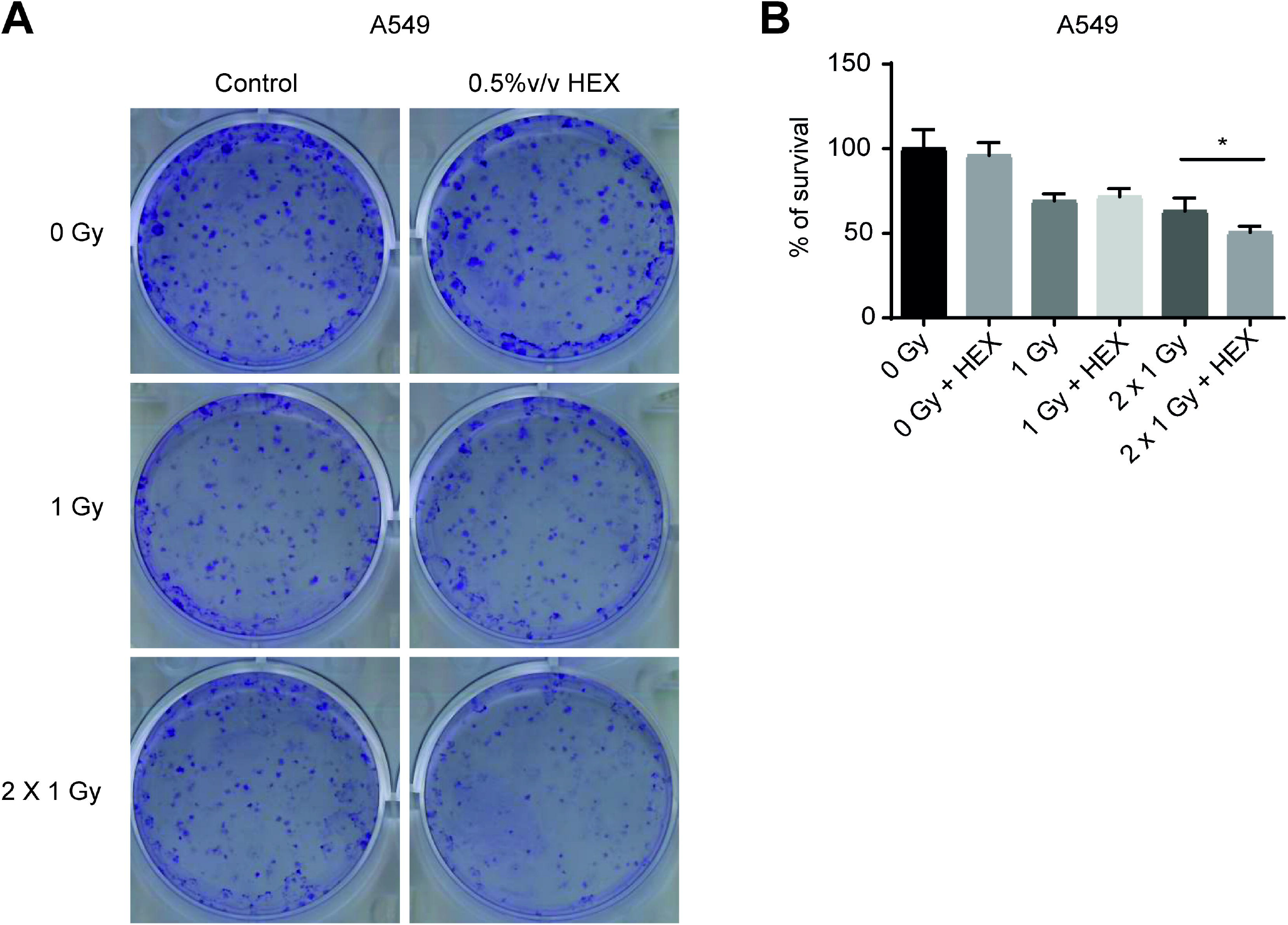
Phase separation is involved in DNA damage response. **A:** 700-700 A549 cells were plated and treated or not with HEX in 0.5 % v/v and exposed or not to 1 Gray (1 Gy) or two times 1 Gy irradiation dose with a 30 min incubation time between the two treatments (2 x 1 Gy). **B:** Chart showing the combined results of the experiment described in A with three independent biological replicates. Survivals are expressed in % of survivals of non-irradiated plates. Statistical significance in the case of comparing 2 x 1 Gy to 2 x 1 Gy+HEX is determined by two-tailed Student’s t-test (*p < 0.05).

## Discussion

In eukaryotes, the two major steps of gene expression are physically separated by compartmentalization. Transcription of chromosomal genes is a nuclear process, whereas translation of the resulting mRNA occurs exclusively in the cytoplasm. Between these two main processes lie numerous steps that provide the opportunity for quality control mechanisms and post-transcriptional tuning of gene expression depending on the demand upon challenges of exogenous origin, such as stress. Compartmentalization by phase separation has recently emerged as a mechanism for post-transcriptional regulation of gene expression, and translation in particular. The formation of phase-separated granules such as P-bodies and SGs, is important for mRNA quality control and stress responses respectively (reviewed in Luo et al., 2018; Decker and Parker, 2012; Guzikowski et al., 2019).

Not1 containing assemblysomes have been described recently as another type of granules that have a role in the co-translational assembly of the proteasome (Panasenko et al., 2019). According to our earlier and current results, the following attributes are important for NCA formation: ribosome pausing during translation, an N-terminally disordered structure, and nascent chain ubiquitylation. Based on these attributes, we used a bioinformatics approach to predict the set of proteins in *Saccharomyces cerevisiae* that might be the subject of NCA formation. For one such candidate protein, Sgs1, we demonstrated that artificially stalled nascent Sgs1 is indeed present in phase-separated granules and, importantly, that the endogenous mRNA is detectable in HEX-sensitive phase-separated granules. This study validated the bioinformatic approach and it is interesting to note that subunits of complexes reported to assemble co-translationally were enriched in the top 350 hits and represent ~15% of our list of 2325 protein complex subunits. For example, Rpt1, Rpt2, Spt20, Taf10 and Set1 are all at the top of the list of our predicted hits (Table S2) and were all reported previously as subunits that are co-translationally assembling with their partner (Panasenko et al., 2019; Kamenova et al., 2019; Halbach et al., 2009; Kassem et al., 2017).

From the following observations, we conclude that NCAs are indeed a type of phase-separated granules that differ from other known granules. First, unlike P-bodies and SGs, NCAs are relatively light assemblies of nascent chain-ribosome complexes and cannot be pelleted by moderate, but only by highspeed centrifugation (Panasenko et al., 2019; Hubstenberger et al., 2017; Jain et al., 2016). Accordingly, Sgs1-containing granules observed by STORM microscopy are smaller than P-bodies or SGs (Van Treeck and Parker, 2019; Nissan and Parker, 2008). Second, NCAs, but not P-bodies and SGs, are cycloheximide- and EDTA-resistant assemblies, and P-bodies, but not NCAs, are RNase-sensitive (Panasenko et al., 2019; Teixeira et al., 2005; Khong et al., 2017). Third, NCAs contain large ribosomal subunits that are absent from SGs (Jain et al., 2016). Finally, NCAs are not formed but actually dismantled upon UV treatment (Figure 2) (Table 1). This observation is particularly exciting, as we will discuss further below. We cannot rule out the possibility that there are transitions from NCAs to, for example, SGs, similar to the communication of P-bodies and SGs (Buchan et al., 2008). Altogether, NCAs are clearly a different type of granular assemblies with different behavior upon stress and therefore having a distinct function.

Our bioinformatic prediction of proteins likely to be translated in NCAs identified components of the DNA damage response. The fact that phase separation is critical for the DNA damage response may provide the clue in understanding the possible role of NCAs. Not only Sgs1, but also other proteins involved in environmental stress responses, including players in the genotoxic stress response, were significantly enriched by bioinformatic predictions among the top 15% of candidates. Stress responses require the production and/or activation of effectors, and this demand for new protein may not be squared due to the transcriptional arrest and defect in translational initiation caused by the stress itself (Holcik and Sonenberg, 2005; Lavigne et al., 2017). However, by storing pre-made stress response mRNAs in the form of RNCs in NCAs, cells can efficiently cope with the situation as such RNCs can continue their translation to produce proteins without the need of *de novo* transcription and/or translational initiation. As combined transcription and translation are relatively time-consuming processes, regulating protein synthesis at a late stage of their production has the potential of fast and timely response to correct, for example, DNA damage even if genes required for the process are themselves affected by the damage, temporarily.

Upon following specific mRNA targets involved in proteotoxic- and others involved in genotoxic stress response we clarified that EDTA-resistant condensates behave very differently under genotoxic and proteotoxic stresses. In the former, they transform into *SGS1, RAD10*, and *RAD14* mRNA into translating ribosomes, and in the latter mRNAs paused during translation of *RPT1* and *RPT2* accumulate in NCAs where they are protected from RQC and await co-translational assembly. Interestingly, NCA to polysome shift is a global response to genotoxic stress and not a targeted one to release mRNAs involved in counteracting genotoxic stress only. In contrast, *RPT1* and *RPT2* mRNAs are specifically increased in EDTA pellets after copper treatment.

We demonstrated that the disorder of the N-terminal region of Sgs1 has a pivotal role in translation regulation since a substitution of two amino acids that changes the disorder propensity of Sgs1, results in RQC and loss of condensation. Therefore, the disordered N-term of Sgs1 is important in keeping expressed nascent Sgs1 protruding from ribosomes in a soluble liquid-liquid phase-separated state. It is important to note that the stress response cannot be managed by the ubiquitous and continuous expression of certain effectors. For example, concerning the context of the DNA damage response, overexpression of Sgs1, a DNA helicase involved in double-strand DNA break repair, is toxic in yeast (Gangloff et al., 1994; Sinclair et al.,1997), or overexpression of the Rad1-Rad10 subunits of Nef1 leads to gross chromosome rearrangements (Hwang et al., 2005). The clear advantage of their delayed completed translation could be that the gene products - required for DNA repair albeit toxic would be expressed in an active form only when the DNA damage appears, avoiding the detrimental effects of unnecessary or excess expression.

Finally, we present evidence that the NCAs characterized in yeast are present also in human A549 cells, as previously shown (Panasenko et al., 2019), and may be relevant also in human cells for the response to DNA damage, thus have a conserved functional role. Indeed, we found that the otherwise radioresistant A549 cell line become more sensitive to DNA damage upon HEX treatment to interfere with phase separation. Bioinformatical analysis to identify NCA-associated proteins in higher eukaryotes may not be as straightforward as in yeast. Indeed, codon usage is an important filter for bioinformatic analysis, and it is defined by tRNA abundance and charging that is known to differ not only between different organisms but within tissues of the same organism. However, our findings here indicate that we can learn more about functional pathways dependent upon NCAs in human cells by extending our analyses in yeast.

## Data availability

The datasets analysed in the current study are available from the corresponding author upon request.

## Acknowledgements

We are grateful for valuable discussions to Dr. Balázs Vedelek and to Dr. Zsuzsa Sarkadi. We thank Jawad Iqbal, Elvira Czvik and Edina Pataki for technical assistance.

## Supplementary figure legends

**Figure S1. Sgs1, Rad10 and Rad14 are strong candidates for being regulated by assemblysomes**. The N-terminal of the proteins shown contain disordered amino acids (blue) and there are pairs of rare codons (encoded amino acids shown with yellow) capable of pausing translation in positions allowing the protrusion of the disordered N-term from the ribosome. Each protein has several lysine amino acids that can be target of ubiquitination in the N-terminal disordered region.

**Figure S2. Ribosomes are EDTA resistant inside granules.**

Non-induced (-Cu) or copper induced (+Cu) yeast cells transformed with copper inducible Sgs1-RNC expressing vectors were extracted and ultracentrifugated through a 60% sucrose cushion in the presence (+) or in the absence (-) of EDTA in concentrations that dissociates the small and large subunits of ribosomes. Total extracts and resolubilized pellets were separated on 10% polyacrylamide gels with electrophoresis and analyzed by Coomassie staining (on the left) and Western blots (on the right) following the Rps6 ribosome subunit. Majority of ribosomal proteins (RP) enriched in the pellet run below 35 kDa as indicated.

**Figure S3. Rpb9 with stop codon is released when expressed from the construct used to generate RNCs for this study.**

**A:** Schematic map of the Rpb9-Released construct. The 122 aa long Rpb9 sequence is N-terminally tagged with a Flag epitope and is under the regulation of copper inducible CUP1 promoter.

**B:** Copper induced Flag-Rpb9 expressing yeast total protein extracts, pellets and supernatants were separated and analyzed as in the case of Sgs1-RNC presented on Fig. 5. As a loading control, we show the Rps3 revelation.

**Figure S4. 1,6-hexanediol dissolves EDTA resistant ribosome assemblies.**

**A:** Same experiment as shown on Figure 2 in the presence or in the absence of 1,6-hexanediol (HEX) in the case samples originating from UV treated yeast cultures. Change in *ACT1, SGS1* and *RAD10* mRNA quantities in EDTA-pellets and total extracts (TE) after indicated treatments as compared to untreated control.

Values in the chart represent average of two independent measurements of biological replicates with error bars representing standard deviation. Statistical significance is determined by Student’s t-test (*p < 0.05, **p <0.01).

**B:** 60% confluent DU145 cell cultures were treated (+HEX) or not with HEX for 30 min, extracted and pelleted in the presence of EDTA with ultracentrifugation through a 60% sucrose cushion. Total extracts (T), top supernatants (SN1), TCA precipitated 60% sucrose cushion (SN2), and pellets (P) were separated on 10% denaturing acrylamide gel and Western blotted to follow Rps6 and Rpb1 proteins with specific antibodies.

**Table S1: Strains, plasmids, oligonucleotides used in this study.**

**Table S2. Yeast protein complex subunits ranked according to their N-terminal disorder probability.** Protein complex subunit IDs are from Swiss-Prot. Ranking was performed after assessment of N-terminal disorder probability exploiting various prediction algorithms. Among these algorithms *vsl2b, iupredLD and espritzXray* was used to generate the final ranking by following the disorder probability on the first 50 and 10 aminoacids (aa’s) respectively. Numbers represent disordered aa’s among the first 10, 25 or 50 aa’s except for the last row where the ubiquitylation prone lysin content of the nascent N-terms are highlighted.

