## Supplementary figures and images for "Phase separated ribosome nascent chain complexes paused in translation are capable to continue expression of proteins playing role in genotoxic stress response upon DNA damage"

### Supplementary Figure 1.

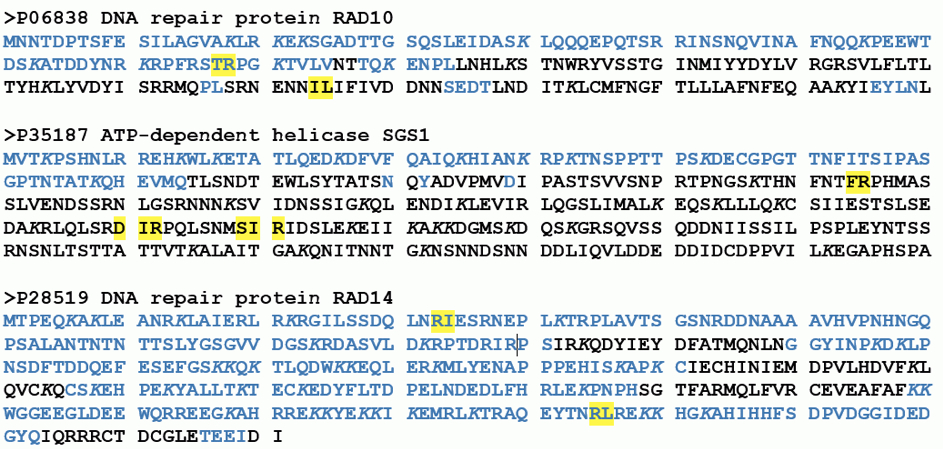

### Supplementary Figure 2.

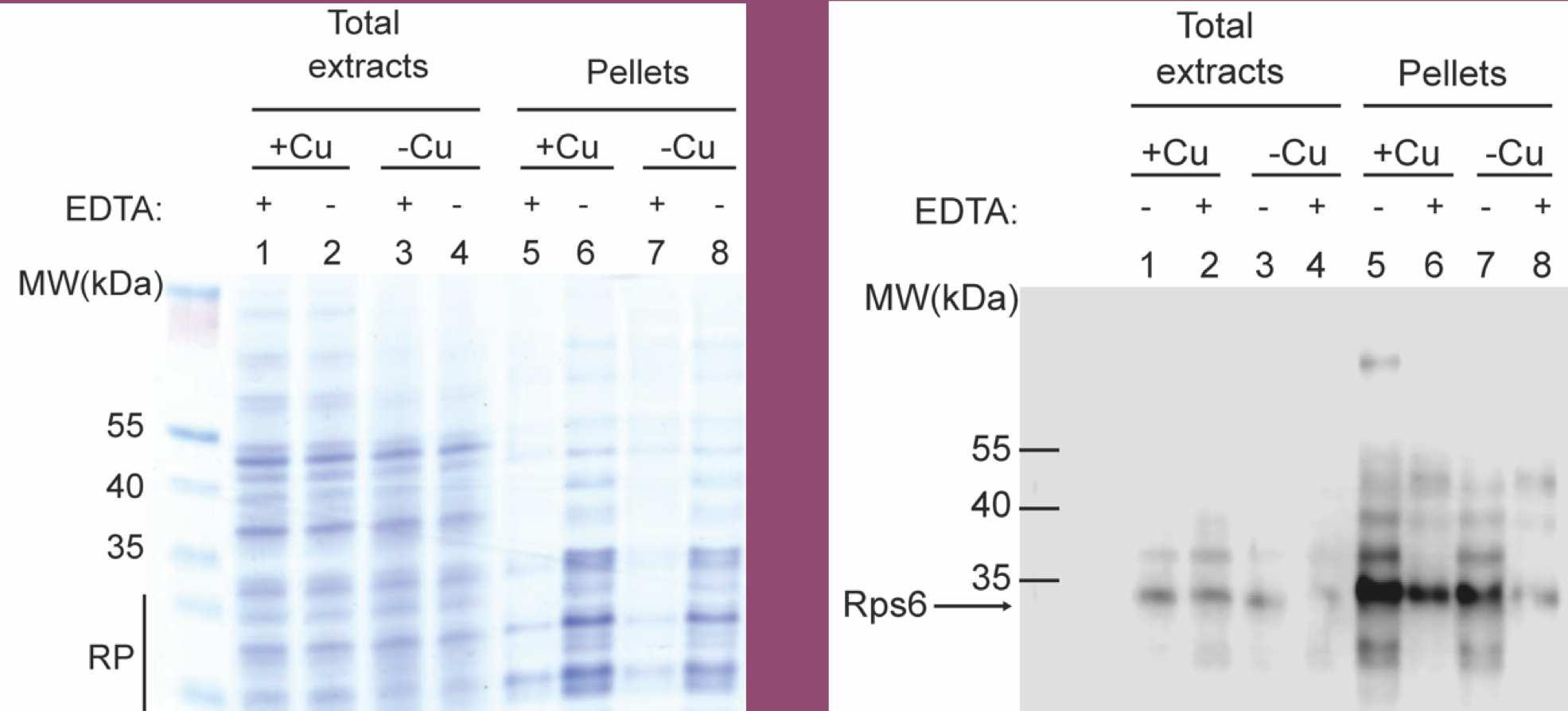

### Supplementary Figure 3.

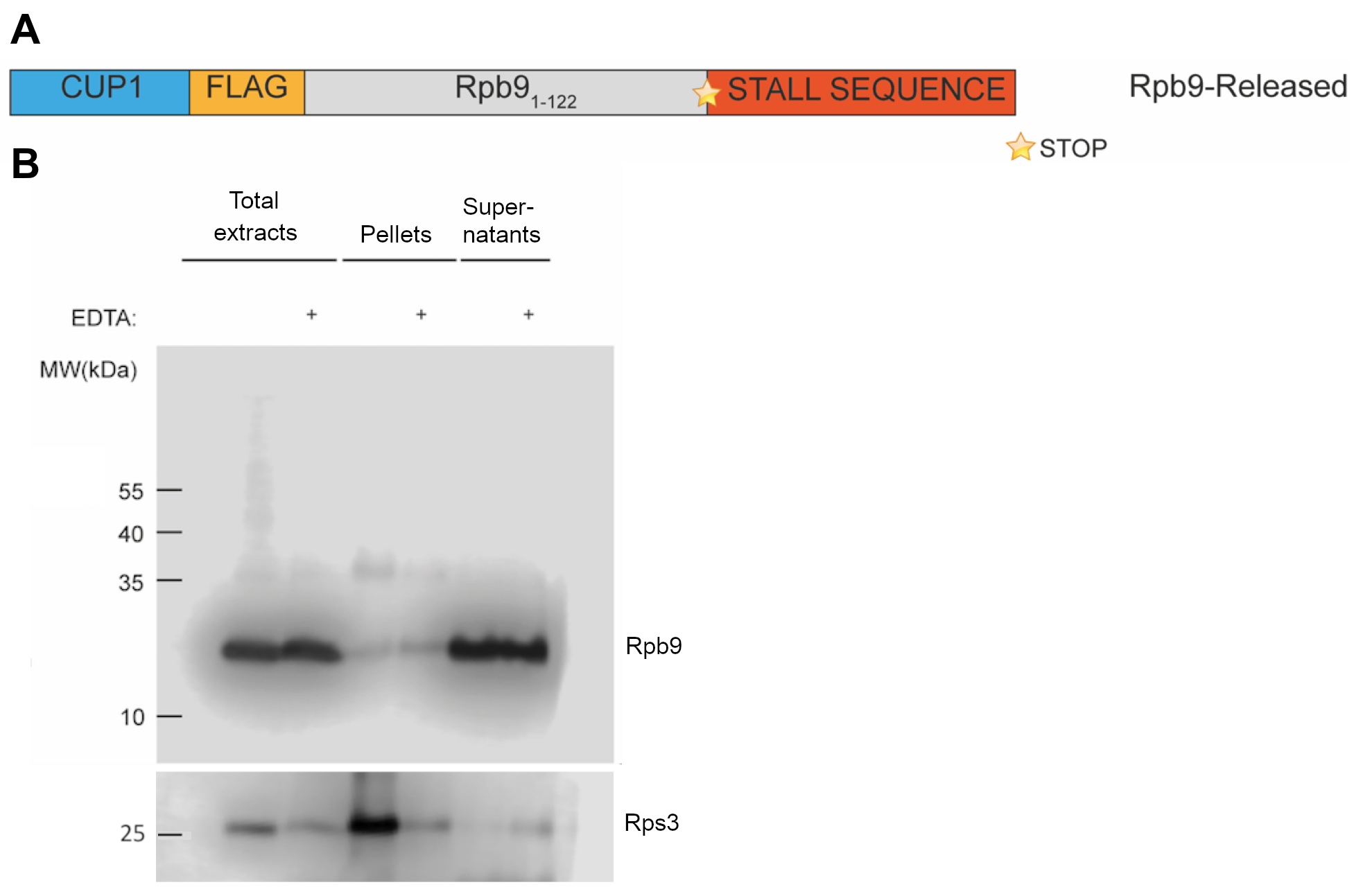

### Supplementary Figure 4.

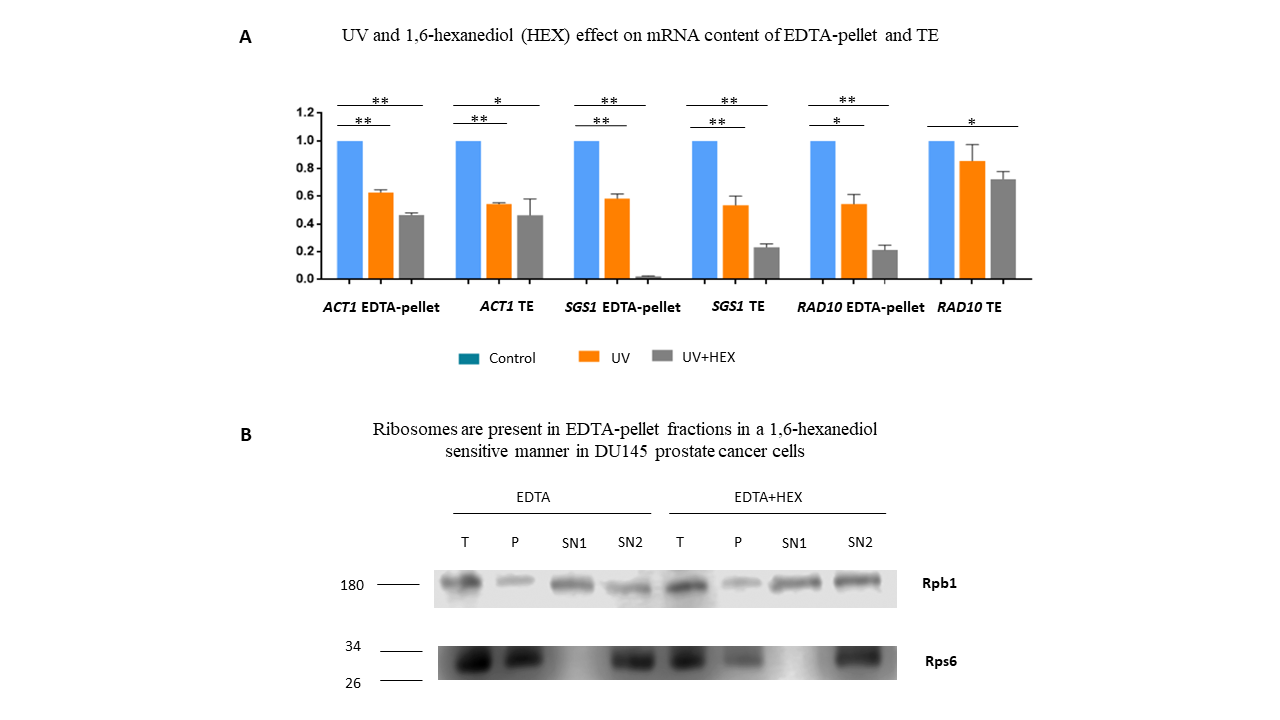
